# Division rate, cell size and proteome allocation: impact on gene expression noise and implications for the dynamics of genetic circuits

**DOI:** 10.1101/209593

**Authors:** François Bertaux, Samuel Marguerat, Vahid Shahrezaei

## Abstract

The cell division rate, size, and gene expression programmes change in response to external conditions. These global changes impact on average concentrations of biomolecule and their variability or noise. Gene expression is inherently stochastic, and noise levels of individual proteins depend on synthesis and degradation rates as well as on cell-cycle dynamics. We have modelled stochastic gene expression inside growing and dividing cells to study the effect of division rates on noise in mRNA and protein expression. We use assumptions and parameters relevant to *Escherichia coli*, for which abundant quantitative data are available. We find that coupling of transcription, but not translation rates to the rate of cell division can result in protein concentration and noise homeostasis across conditions. Interestingly, we find that the increased cell size at fast division rates, observed in *E. coli* d other unicellular organisms, buffers noise levels even for proteins with decreased expression at faster growth. We then investigate the functional importance of these regulations using gene regulatory networks that exhibit bi-stability and oscillations. We find that network topology affects robustness to changes in division rate in complex and unexpected ways. In particular, a simple model of persistence, based on global physiological feedback, predicts increased proportion of persistors cells at slow division rates. Altogether, our study reveals how cell size regulation in response to cell division rate could help controlling gene expression noise. It also highlights that understanding of circuits’ robustness across growth conditions is key for the effective design of synthetic biological systems.

## Introduction

Microbial species can proliferate in a variety of environmental conditions. How genomes achieve this phenotypic flexibility is a fundamental biological question. Regulated gene expression is a key mechanism by which cells adapt physiologically to changing environments. For example, different types of metabolic enzymes are expressed to support growth on different carbon sources (Görke & Stülke, 2008). Despite this remarkable adaptability, the rate at which cells proliferate can vary strongly from one environment to another. For example, *E. coli* division rates range between 0.5 to 3.5 doublings per hour in response to different carbon sources (Taheri-Araghi *et al*, 2015).

In addition to specific gene regulation, changes in division rate are accompanied by global physiological changes (Figure 1), such as changes in cell size at division and gene expression. Global changes in gene expression with cellular growth rates are required to counteract the increase in dilution rate inherent to faster proliferation and maintain average protein concentrations. This global coordination of gene expression with the division rate could involve changes in transcription, translation and mRNA turnover. Experimental evidence suggests that in yeast and bacteria this coordination occurs primarily at the level of transcription (Keren *et al*, 2013; Gerosa *et al*, 2013; Berthoumieux *et al*, 2013; García-Martínez *et al*, 2016). Consistent with this, bacteria global translation rates are less affected by the rate of division than transcription rates, except at very slow division rates (Klumpp *et al*, 2013; Dai *et al*, 2016). In *B. subtilis*, the translation rate per mRNA has even been found to decrease, while the total mRNA concentration doubles as the division rate doubles (Borkowski *et al*, 2016). Messenger RNA degradation rates as well remain largely unchanged between slow and fast growth in *E. coli* (Bernstein *et al*, 2002). In yeast, however, mRNA degradation rates have been proposed to be globally regulated by the division rate (García-Martínez *et al*, 2016). Together, these observations suggest that coupling of either transcriptional or post-transcriptional layers of regulation with the division rate can result in protein expression homeostasis. However, it remains unclear whether coupling of trancription with growth rates, rather than mRNA degradation or translation for instance, has a particular impact on protein expression dynamics, variability, or even the cell fitness.

**Figure 1:**
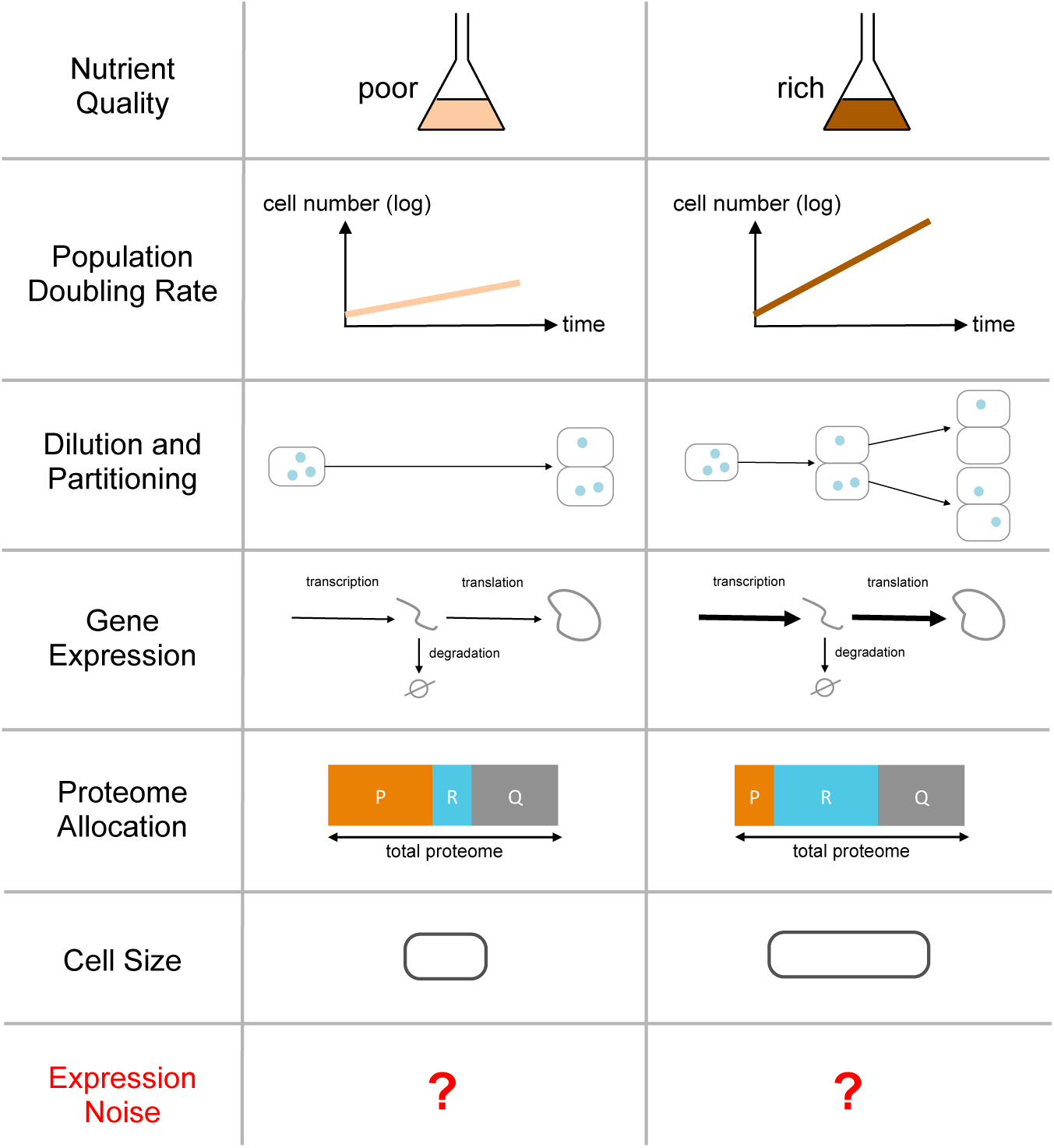
Global cellular factors affecting gene expression noise that depend on growth conditions. Nutrient quality can increase the population doubling rate by promoting growth and division of individual cells. This leads to increased dilution of molecules, and more frequent random partitioning of molecules between daughter cells. Because faster growth requires a higher rate of cell mass production, rates of mRNA and protein expression increase globally with the division rate. However, the relative changes in mRNA and protein expression rates is gene-dependent because the proteome composition is reshaped when the division rate changes (Scott *et al*, 2014). For example, the fraction of ribosomal proteins (*R* proteins) will increase with the division rate while the fraction of metabolic enzymes (and other *P* proteins) will decrease, the fraction of house keeping proteins (and other *Q* proteins) remain constant (Scott *et al*, 2010). Cell size as well is known to increase with the division rate in response to nutrient-based modulations (Schaechter *et al*, 1958; Basan *et al*, 2015). All those factors affect both average expression and expression noise in a non-trivial manner.

The dependence of gene expression parameters on the division rate has been shown to vary between genes. Specifically, the fraction of the proteome occupied by genes from different functional classes has been shown to vary with the division rate following specific and simple trends (Scott *et al*, 2010; Li *et al*, 2014; Hui *et al*, 2015). For a given type of division rate modulation, proteins can be categorised in three classes called *R, P* and *Q* depending on whether their proteome fraction respectively increases, decreases or is maintained with the division rate respectively. Simple models of proteome allocation and cell physiology have shown that the changes in global protein fractions observed experimentally are consistent with a maximisation of the division rate (Molenaar *et al*, 2009; Scott *et al*, 2014; Goelzer & Fromion, 2017). For example, ribosomal proteins that constitute most of the *R* proteins are needed in larger amounts to support fast growth in rich media (Scott *et al*, 2010). As a consequence of a large *R* sector at fast growth another group of proteins, such as transporters and metabolic enzymes for instance, decrease in concentration (class *P*), as proteome fractions add up to one. Finally, the class called *Q* contains housekeeping proteins, whose proteome fraction is maintained across all conditions.

Because total protein concentration is approximately constant across conditions (Basan *et al*, 2015), the concentration of *P* proteins decreases at fast growth. Lower concentrations mean lower number of molecules per unit of volume. Intrinsic noise of *P* proteins, which results from the random timing of biochemical reactions and depends on absolute molecule numbers rather than concentrations, could therefore be higher at fast growth. Intrinsic noise contributes to cell-to-cell variability in gene expression and leads to non-genetic phenotypic variability (Shahrezaei & Swain, 2008). In addition, gene expression is affected by *other* stochastic and dynamic cellular processes, resulting in so-called extrinsic noise (Elowitz *et al*, 2002; Shahrezaei *et al*, 2008). An important source of extrinsic noise in gene expression stems from the processes associated with the cell cycle, including cell growth and cell division, as illustrated by the experimental and modelling studies discussed below. Mathematical modelling has suggested that random partitioning of biomolecules at cell division is an important source of noise in gene expression and hard to separate from intrinsic noise (Huh & Paulsson, 2011). Other modelling studies have highlighted the contribution of heterogeneity in cell cycle time on noise in gene expression (Johnston *et al*, 2012; Schwabe & Bruggeman, 2014; Antunes & Singh, 2014; Soltani *et al*, 2016). Moreover, cell cycle dependent expression and the timing of DNA replication also influences noise in gene expression in unexpected ways (Luo *et al*, 2013; Schwabe & Bruggeman, 2014; Peterson *et al*, 2015; Soltani *et al*, 2016). Several experimental studies have identified the cell cycle as a major source of noise in gene expression in bacteria and yeast (Cookson *et al*, 2010; Zopf *et al*, 2013; Keren *et al*, 2015; Walker *et al*, 2016). These studies suggest that gene expression noise is generally higher at lower division rates (Keren *et al*, 2015; Walker *et al*, 2016). The impact of cell division and random partitioning of molecules on the behaviour of simple circuits has also been studied by modelling (Gonze, 2013; Lloyd-Price *et al*, 2014; Bierbaum & Klumpp, 2015). It has been shown that simple genetic oscillators can sustain oscillation in the presence of cell division but the oscillations could be entrained by the cell cycle depending on the circuit topology (Gonze, 2013). Random partitioning of biomolecules at division can also affect the dynamics of simple circuits such as the stability of biological switches for instance (Lloyd-Price *et al*, 2014).

Cell size is regulated both during the division cycles and between different growth conditions. Although being a long-standing problem in cell biology, the mechanisms behind cell size homeostasis remain largely elusive. Interest for this question has been recently renewed, particularly in bacteria. Recent data suggests that many bacterial species follow a so-called adder principle of growth, adding a constant cytoplasm volume in each division cycle, independently of their size at birth. As DNA replication is controlled by size in *E. coli*, understanding whether this phenomenon is participating to cell size homeostasis and the adder principle across growth conditions is currently the focus of intense investigation both at the theoretical and experimental level (Ho & Amir, 2015; Wallden *et al*, 2016; Si *et al*, 2017). Interestingly, cell size at division is positively correlated with division rates in both bacteria and yeast, cells becoming larger in richer environments (Schaechter *et al*, 1958; Turner *et al*, 2012). This phenomenon is universal across unicellular organisms but there is no satisfying explanation of why regulation of cell size with growth conditions has evolved.

Global regulation of gene expression and cell size is likely to affect the dynamics and function of genetic and biochemical networks inside cells (Shahrezaei & Marguerat, 2015). A pioneering study quantified how division rate dependent global regulation of gene expression affects the average concentration of a constitutively expressed gene product, and how this in turn can affect the behaviour of simple synthetic genetic networks (Klumpp *et al*, 2009). Another theoretical study showed that the division rate dependence of gene expression could impact the qualitative behaviour of a synthetic oscillator circuit, the ‘repressilator’ (Osella & Lagomarsino, 2013). Moreover, the division rate regulation of a gene impacting fitness can result in non-trivial global feedback in gene regulation (Klumpp *et al*, 2009; Kiviet *et al*, 2014; Tan *et al*, 2009). However, a comprehensive view of how global changes in gene expression and cell size in response to growth conditions impact on gene expression noise is lacking.

In this study, we shed light on the regulation of noise in gene expression across growth conditions in the bacterium *E. coli* by integrating existing data on global regulation of gene expression and cell size into detailed computational models of stochastic gene expression in growing and dividing cells. We then use examples of some simple genetic networks to illustrate how the changes in gene expression noise across growth conditions affects the dynamics of cellular systems.

## Results

### Stochastic gene expression in growing and dividing cells

To fully capture the effect of cell cycle on noise in gene expression, we model the stochastic expression of a single gene in growing and dividing cells (Figure 2 A-B, Supplemental Figure 1-A). Transcription, mRNA degradation and translation are represented by single stochastic reactions. Corresponding rates are noted *k*_*m*_, *γ*_*m*_ and *k*_*p*_ respectively. Because the majority of *E. coli* proteins are stable (Goldberg & St John, 1976), we first neglect protein degradation. During the cell cycle, we assume cell size increases exponentially at a fixed rate that results in a decrease in the concentration of the mRNA and the protein when their numbers of molecules do not change. We model cell division as a discrete event that splits the cell volume in two, and each molecule is randomly partitioned between daughter cells with a probability matching their inherited volume fraction. In our simulations, we keep only one of the two daughter cells, therefore reproducing the popular *mother machine* experimental setting (Wang *et al*, 2010). Our general simulation algorithm is described in details in the Methods section *Simulation algorithm*.

**Figure 2:**
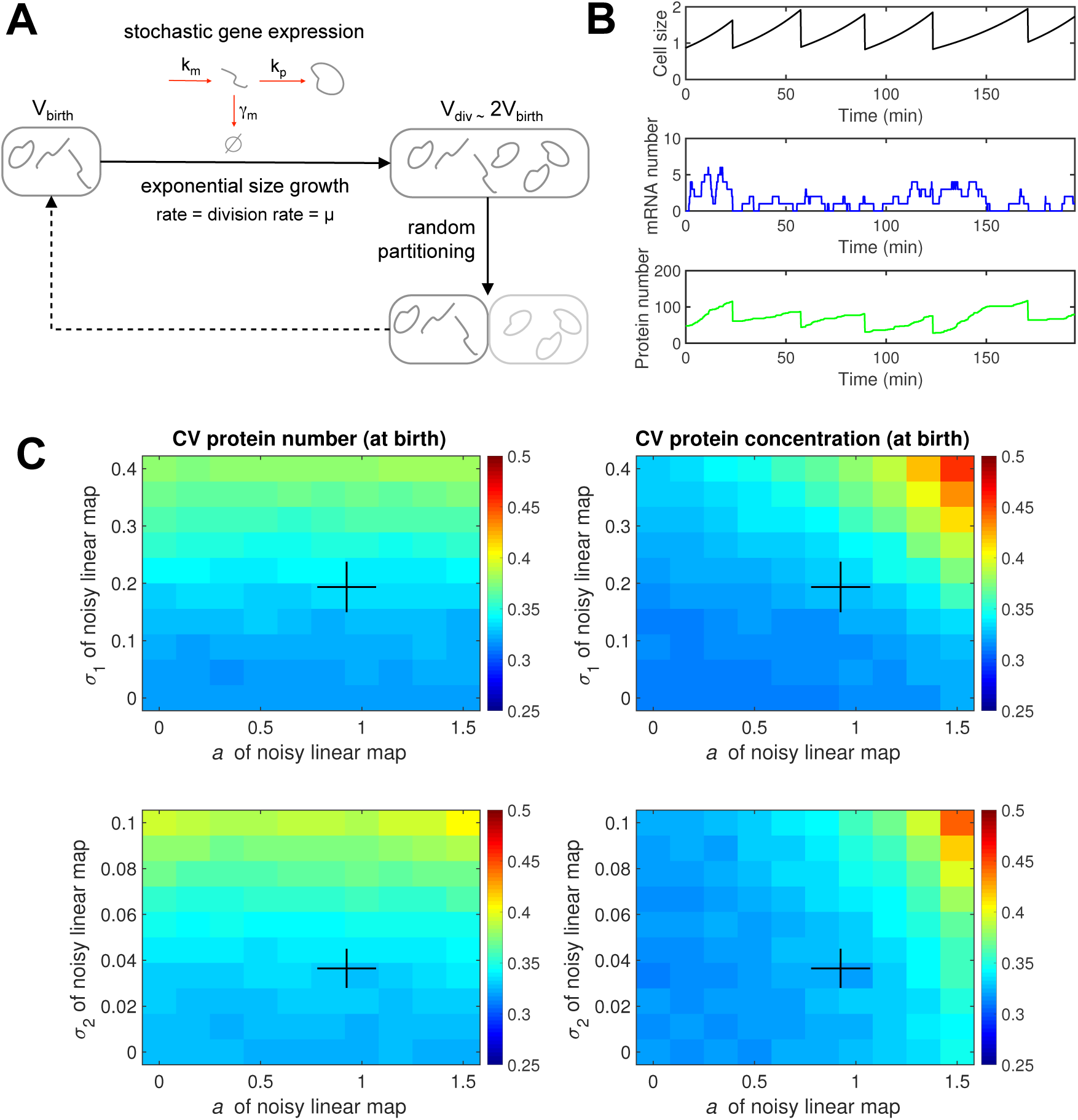
Modelling stochastic gene expression in growing and dividing cells. **(A)** Sketch of the modelling approach. See Methods for details. (B) Example of simulated trajectories. Typical parameters for *E. coli* have been used (See Methods). (C) Impact of noisy linear map (NLM, See Methods) parameters on protein noise. Heatmaps of protein number noise (left) or concentration noise (right) (defined as the coefficient of variation, CV, across newly born cells) when *a* and *σ*_1_ (top) or *a* and *σ*_2_ (right) are varied. Other parameters are kept constant at reference values, except *b* that changes with *a* such that the average size at birth is constant. Black crosses indicate empirical ranges estimated from mother machine data (See Methods and Supplemental Figure 1).

Cellular growth rate, cell size at division, and cell size at birth are known to vary between individual cells even in identical, tightly controlled conditions. Variability in size at birth arises from variability in the mother cell size at division but also from imperfect volume splitting between the two daughter cells. To realistically account for this variability, we use the *noisy linear map* (NLM) model (See Methods and Supplemental Figure 1), a recent phenomenological model of cell size control that captures the variability in cell size at birth and division observed experimentally as well as their correlation within individual cell cycles (Tanouchi *et al*, 2015; Jun & Taheri-Araghi, 2015; see also Amir, 2014 for a closely related approach). The degree of this correlation is related to the mechanisms underlying cell size homeostasis. For example, a noisy linear map with the parameter *a* equal to 1 corresponds to an adder strategy, where a fixed cytoplasm volume is added to the cell between each division. Alternatively, a parameter *a* equal to zero corresponds to a sizer strategy, where cell division is triggered at a fixed size (Jun & Taheri-Araghi, 2015). Noise in the NLM model is represented via two parameters, *σ*_1_ and *σ*_2_, accounting respectively for the noise in division size of the mother cell and the noise in birth size that results from imperfect splitting of the mother.

We have first analysed the effect of the NLM parameters on the gene expression noise for an intermediate growth rate. In Figure 2-C, we show protein number and concentration noise (CV) at cell birth immediately after cell division and at the beginning of the cell cycle when the noise in division size (*σ*_1_), noise in size splitting (*σ*_2_) and *a* are varied. When noise in the NLM is large (*σ*_1_ or *σ*_2_), we observe increased noise in protein number at the beginning of the cell cycle (Figure 2-C). This is due to partitioning noise, as this increase is almost invisible at the middle of the cell cycle (Supplemental Figure 2). Moreover, we find that protein concentration noise is only marginally affected by NLM noise, as we assume probability of random partitioning of biomolecules is proportional to the inherited volume of the daughter cells after division. This result is consistent with previous work (Schwabe & Bruggeman, 2014) and extends it by explicitly modelling both mRNA and protein molecules. For values of *a* greater than one size control is not very effective in filtering noise in cell size and there is an increased size variability for large *a* and large NLM noise (*σ*_1_ or *σ*_2_), as also found by others (Modi *et al*, 2017). As a result the protein concentration noise that directly depends on cell volume shows an increase at large *a* and large NLM noise.

A priori, it is possible that the NLM parameters that best describe a given single-cell dataset could change with growth conditions. Therefore, we have inferred the parameters of the NLM from a recent mother machine dataset of cells grown in 7 different carbon sources supporting a wide range of division rates (Taheri-Araghi *et al*, 2015). We find that NLM parameters can indeed change with the division rate (Supplemental Figure 1). Notably, the slope parameter *a* is significantly lower than 1 at slow growth, consistent with another study reporting a deviation towards a sizer strategy (*a* < 1) in slow regimes (Wallden *et al*, 2016). Modulation of *a* has also been reported for other bacteria (Priestman *et al*, 2017). In addition, individual cell growth rates are well described by normal distributions in all conditions. Based on that analysis, we derive linear functions describing NLM parameters as a function of the division rate (Supplemental Figure 1). This enables us to realistically model growth and division at the single cell level over a wide range of division rates and investigate their effects on gene expression noise.

### Expression noise depends on division rate even when protein concentration is maintained

We consider first genes whose protein concentration stays constant when the division rate changes (*Q* class proteins). Interestingly, this expression strategy requires that at least one of the gene expression rates *k*_*m*_ (transcription rate), *γ*_*m*_ (mRNA degradation rate) or *k*_*p*_ (translation rate per mRNA) changes with the division rate to compensate for increased dilution of mRNA and protein molecules.

Using our model and typical values for gene expression rates at 2 doublings per hour as a baseline, we computed the change in protein concentration *noise* with the division rate when average concentration is maintained either by adapting the transcription rate only (Figure 3-A) or the translation rate per mRNA only (Figure 3-B). To investigate the contribution of distinct sources of noise and of variability in cell size to protein concentration noise we consider multiple scenarios in which different sources of variability are turned off (colour codes in Figures 3-A and 3-B).

**Figure 3:**
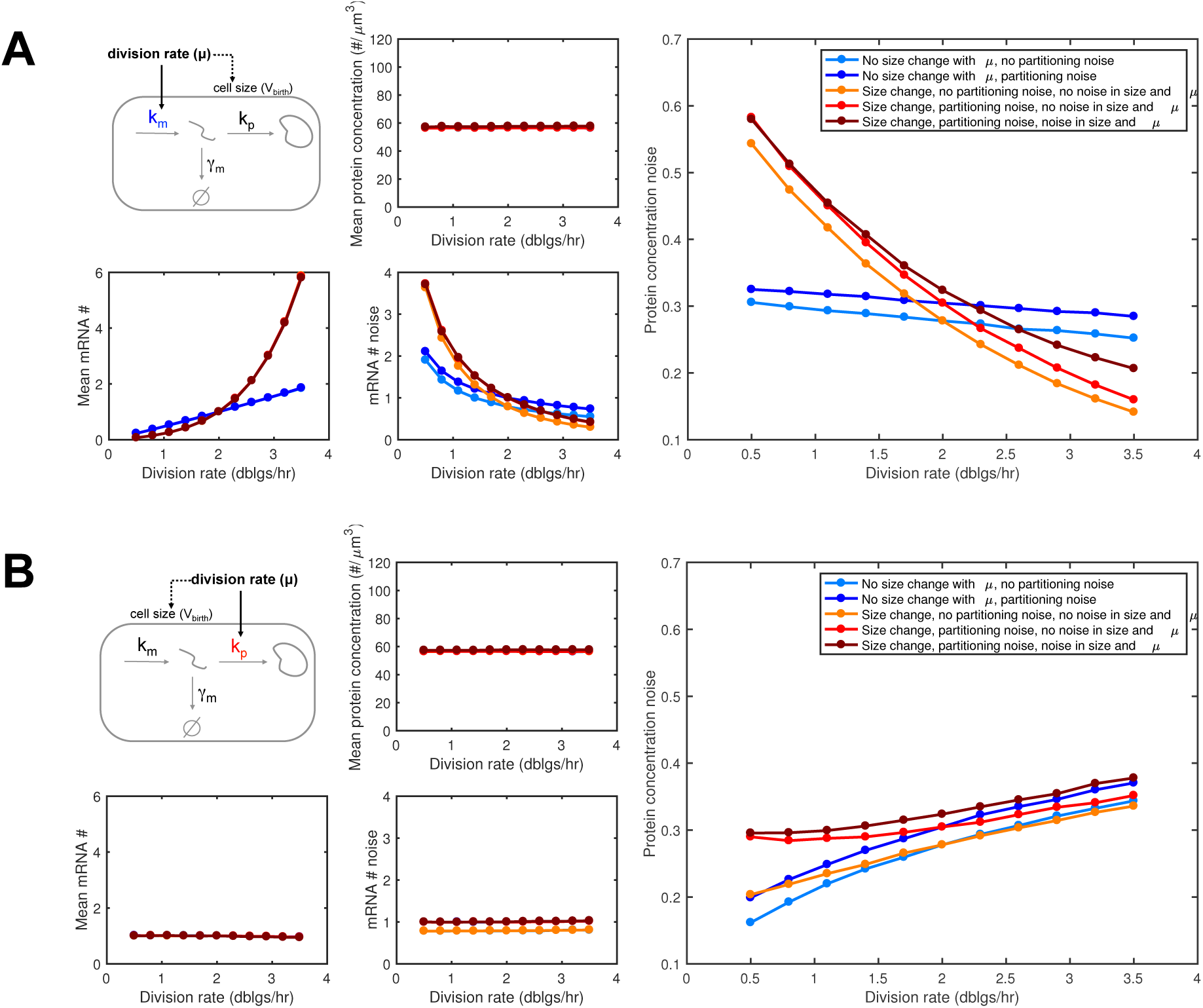
Changes in cell size, transcription and translation rates with the division rate impact expression noise even when average protein concentration is maintained *(Q* expression). **(A)** Change of protein concentration noise (right) with division rate when the average concentration is maintained (middle-top plot) by tuning the transcription rate (left-top plot). Noise is the CV of protein concentration across newly born cells. The mRNA average number *(Jf)* and CV in are also shown (bottom-left plots). Different model variants are simulated to explore the contribution of random partitioning noise, size change with the division rate, and noise in size (NLM parameters) and cellular growth rate (See Methods and Supplemental Figure f). (B) Same as (A) but when the translation rate is tuned instead of the transcription rate. (A **and** B) The corresponding gene expression rates dependency with the division rate are shown in Supplemental Figure 4.

Our simulation results reveal that maintaining average protein concentration of *Q* proteins by adjusting transcription or translation to the division rate leads to very different behaviours of the protein concentration noise. Moreover, we find that the increase of cell size with division rate observed across unicellular organisms strongly contributes to these behaviours. In the case of transcription rate adjustment, protein noise sharply decreases with the division rate. A milder decrease is also observed when cell size is kept constant across division rates. In the case of translation rate adjustment, protein noise increases with the division rate instead, whether cell size changes or not.

To better understand these results, we looked at how mRNA numbers change with the division rate in the different situations (bottom left plots in Figures 3-A and 3-B). When transcription adjusts to the division rate in order to maintain average protein expression, mRNA numbers increase with the division rate. As mRNA noise (mRNA numbers are typically much lower than protein numbers) is a major contributor to protein noise, an increase in mRNA numbers results in a decrease in protein noise. However, when translation adjusts to the division rate instead, mRNA numbers remain mostly unchanged. This is possible, because mRNA degradation rates are large compared to the division rate, resulting in mRNA numbers being less sensitive to dilution than protein numbers. Despite little change in mRNA numbers and hence mRNA noise, the increase in protein noise can be explained by a higher propagation of the mRNA noise to protein, since contribution of transcription to protein noise depends on the ratio of mRNA lifetime, which is mostly constant, and protein lifetime, which is set by the dilution rate (itself set by the division rate, Swain *et al*, 2002).

While the relative contribution of stochastic gene expression, partitioning noise, variability in cell growth rate, cell division size and cell birth size to total protein noise can change with the division rate, we find that the contribution of stochastic gene expression is the major source of noise at all division rates (Supplemental Figure 3). Other sources of noise show only minor contributions to overall noise levels. Namely we observe a small contribution of partitioning noise to total noise that remains constant or decreases with growth when transcription or translation are adjusted to division rate respectively. NLM noise showed an opposite trend with a small contribution to total noise increasing at fast division rates when transcription is adjusted and a constant contribution to noise in the case of coupling to translation (Supplemental Figure 3). In summary, our simulations demonstrate that for genes with typical expression parameters at intermediate division rates, maintaining a constant protein concentration across growth conditions by adjusting transcription to the division rate leads to a decrease of protein noise. In contrast, adjusting translation to the division rate increases protein noise levels. Moreover we find that the main source of noise (with the parameters used) is the intrinsic stochasticity in the gene expression reactions.

### Increase of cell size with the division rate prevents noise increase for consti-tutively expressed proteins despite a decrease in average concentration

The results described above concern proteins belonging to the *Q* category, whose average concentration is maintained constant independently of the division rate. Klumpp and colleagues have shown that constitutively expressed proteins instead belong to the *P* category: their concentration is decreased at fast growth (Klumpp *et al*, 2009). The transcription rate of constitutively expressed genes strongly increases with the division rate, while mRNA degradation rate and translation rate per mRNA remain relatively constant (Klumpp *et al*, 2009). However, this is not sufficient to balance both increased dilution and increased cell size (Klumpp *et al* (2009), Supplemental Figure 4 and Figure 4 top left plot).

**Figure 4:**
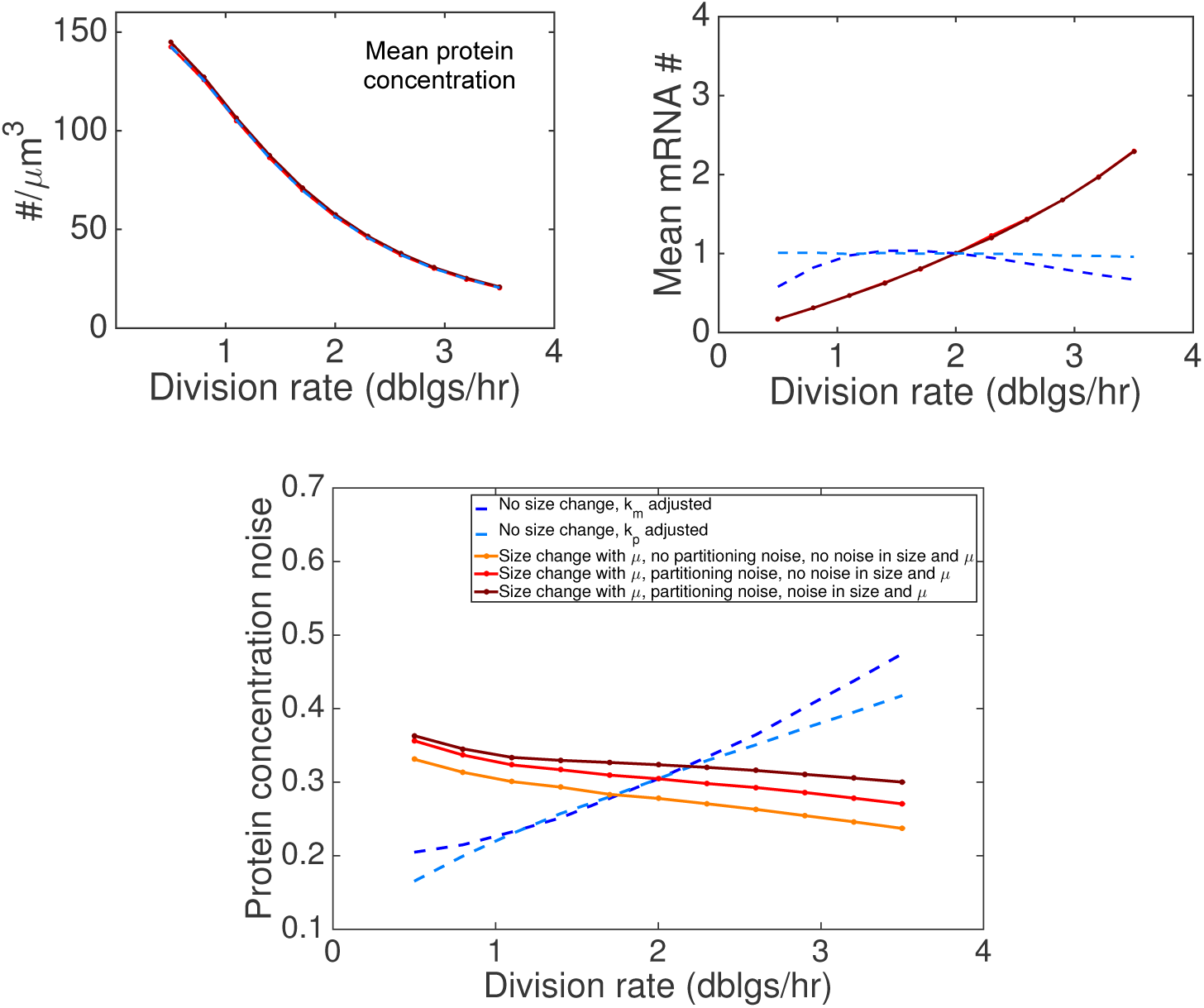
Larger cell size at fast division rates prevents expression noise increase despite a decrease in average concentration (*P* expression). To reproduce *P* expression, we used gene expression parameter dependencies with division rate for constitutively expressed proteins extracted from a previous study ((Klumpp *et al*, 2009), See Methods and Supplemental Figure 4 for details). Average protein concentration (top left), average mRNA number (#) (top right) and protein concentration noise (bottom) are shown. The same model variants as in Figure 3 were used. Two additional scenarios are also shown, in which cell size does not change with division rate but either the transcription rate (dashed dark blue) or the translation rate (dashed light blue) is adjusted to obtain the same decrease of average protein concentration with division rate (other parameters remaining constant and equal to the reference values of solid line simulations at 2 doublings per hour).

Remarkably, using parameters of gene expression from (Klumpp *et al*, 2009) (See Methods and Supplemental Figure 4), we find that protein noise decreases with division rate, despite a strong decrease in average protein concentration (Figure 4). Cell size increase with division rate is a key contributor to this behaviour. Assuming that increased expression noise for *P* proteins at fast growth is deleterious, this observation could explain why increased cell size at fast division rates is a universally conserved feature of unicellular organisms.

As for the *Q* protein described above, we find that the relative contribution contribution of intrinsic stochastic gene expression is predominant at all division rates for *P* proteins (Supplemental Figure 3). However, contribution of both partitioning noise and size and growth rate variability does also increase, but moderately, at fast division rates.

So far, we have focused on gene with low baseline expression and high intrinsic noise. In Supplemental Figure 5, we show that for *P*-like proteins with increasing baseline expression levels (through changes in transcription and/or translation rates), the predicted decrease of protein concentration noise with the growth rate gets weaker, eventually reaching a zero intrinsic noise limit, which is approximately independent of the division rate.

**Figure 5:**
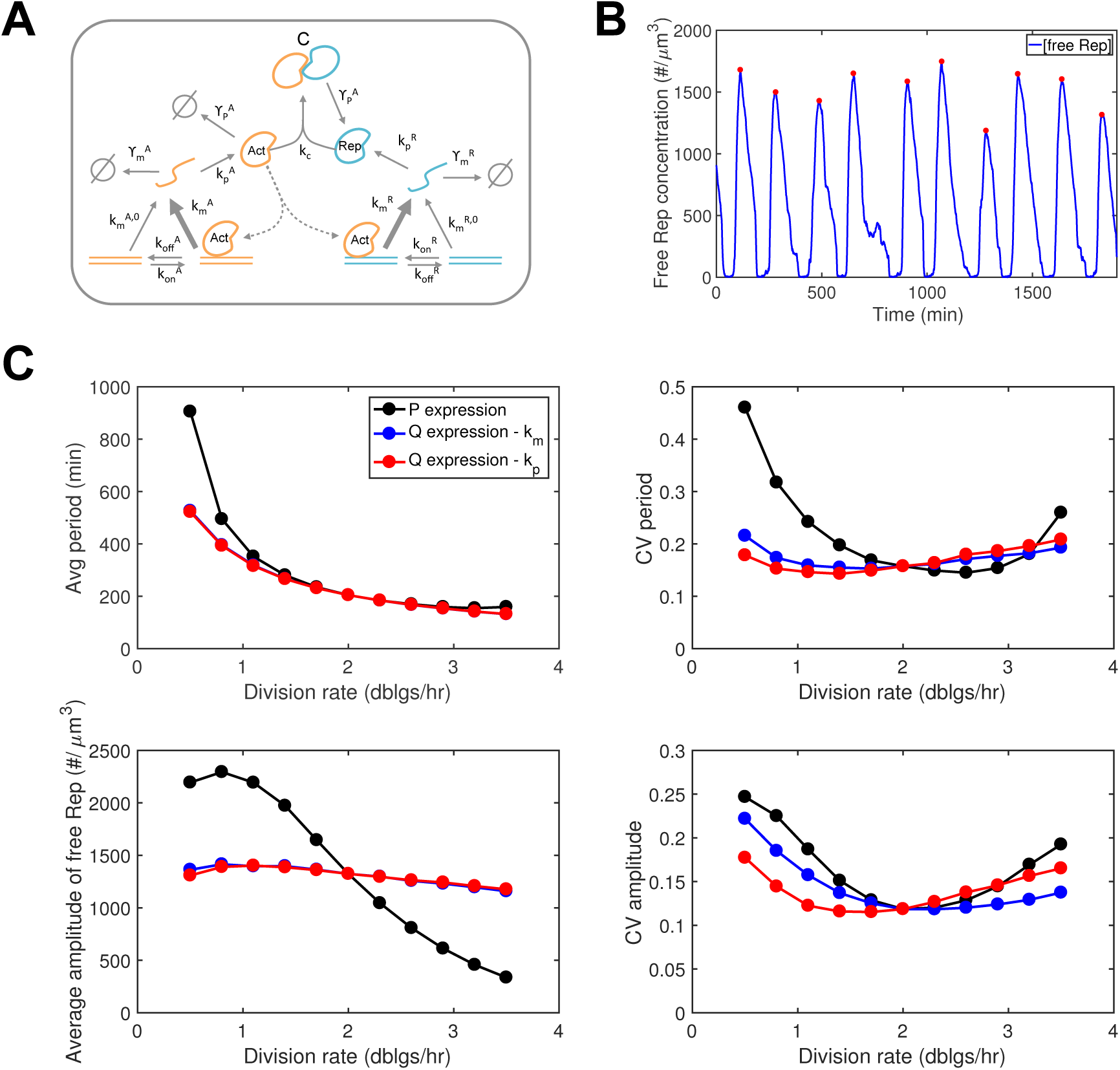
Behaviour of an oscillator circuit at different division rates. **(A)** Schematic of the oscillator circuit described in (Vilar *et al*, 2002). See Methods for model description and parameter values. (B) Example simulation showing oscillations in free *Rep* concentration. Detected peaks are shown with red circles. Note that the timescale of oscillations is around 3 hours, while the inter-division time is around 30 minutes. (C) Change of the oscillatory behavior (average period, noise in period, average amplitude, noise in amplitude) as a function of division rate. The black curves correspond to *P* expression. The other curves correspond to situations in which either transcription rates (blue) or translation rates (red) are increasing with division rate in order to maintain average expression (*Q* expression in absence of binding of *Act* with *Rep*).

### Impact of division rate on the behaviour of an oscillator circuit

Changes in average expression and noise of individual proteins with the division rate in response to environmental changes is likely to impact the behaviour of genetic circuits (Klumpp *et al*, 2009). Even when the protein average expression (in isolation, i.e. without the circuit-specific regulations) is maintained, the expression noise can still change (Figure 2) meaning that circuit behaviour could depend on the division rate (Shahrezaei & Marguerat, 2015).

To investigate these effects, we first consider a canonical two proteins oscillator circuit recapitulating essential features of natural clocks (Figure 5-A) (Vilar *et al*, 2002). An actively degraded activator protein (*Act*) promotes its own transcription as well as the transcription of a stable repressor protein (*Rep*) by promoter binding. *Rep* can also bind *Act*, preventing it to bind promoters. This circuit can lead to oscillations as illustrated in Figure 5-B. A detailed analysis of why oscillations arise is beyond the scope of this study and has been explored before (Guantes & Poyatos, 2006; Kut *et al*, 2009). Briefly, because *Rep* competes with promoters for the binding of *Act*, when the amount of free *Rep* is large only basal transcriptional activity for both genes is possible. Because *Rep* is stable, such a state can last until dilution and partitioning renders free *Rep* levels too low to efficiently prevent promoter binding by *Act*. Promoter activation leads to a burst of *Act* by auto-activation, but *Rep* levels eventually rise because *Act* also promotes *Rep* transcription. When *Rep* levels are sufficient to efficiently compete with *Act* promoter binding, a novel cycle starts.

We asked how the circuit behaviour was affected when division rate modified. We first assume that basal transcription, translation and mRNA degradation follows the same dependency as constitutively expressed proteins (i.e. *P* proteins, as in Figure 4), and that the fold-change increase of transcription rate when the promoter is activated by *Act* is independent of the division rate. The resulting changes in circuit behaviour with the division rate are shown in Figure 5-C (black lines). The average period increases as the division rate decreases because dilution is an important driver of the oscillations. The average amplitude of free *Rep* oscillations is also strongly dependent on the division rate, and decreases as the division rate increases. This is consistent with *P* expression, although different behaviours are in theory possible because of gene regulation. The noise in circuit behaviour changes as well with the division rate. Specifically, noise in period and amplitude of the oscillations display ‘U’ shape dependencies with the division rate, with lower noise close to the reference division rate of 2 doublings per hour. In summary, constitutive expression (typical of *P* proteins) leads to changes in average behaviour and a strong increase in noise of an oscillatory circuit at very low or very high division rates.

We then investigated whether *Q* expression of the circuit components could increase the robustness of oscillations in response to changes in division rate. As in Figure 3 we consider two modes of *Q* expression, either by transcriptional adjustment (blue) or translational adjustment (red). Both modes could maintain the average amplitude of oscillations in a narrow range, but the average period remained strongly dependent on the division rate (Figure 5-C). While both modes resulted in identical changes in circuit average behaviour, they led to slightly different dependencies of noise in oscillations with the division rate. The division rate with the minimal noise in amplitude is around 2.3 doublings per hour for transcriptional adjustment and around 1.5 doublings per hour for translational adjustment. In summary, *Q* expression increased robustness of oscillations compared to constitutive (*P*) expression, but it is not sufficient to make the oscillator’s period independent of the division rate. *Q* expression via transcriptional or translational adjustment led to similar, but not identical changes of noise in oscillations with the division rate.

### Impact of division rate on the behaviour of the toggle switch

We investigate next a simple synthetic circuit known to exhibit bistability: the toggle switch (Gardner *et al*, 2000), in which two proteins repress each other’s transcription (Figure 6-A). We asked first whether different circuit behaviours, namely the existence of bistability, the occupancy of the states, and the switching rates between states, were affected by changes in division rate and adjustment of transcription or translation to division rates. To this end, we consider simple model assumptions that are sufficient to generate stochastic switching between different states (Methods) with typical parameter values.

**Figure 6:**
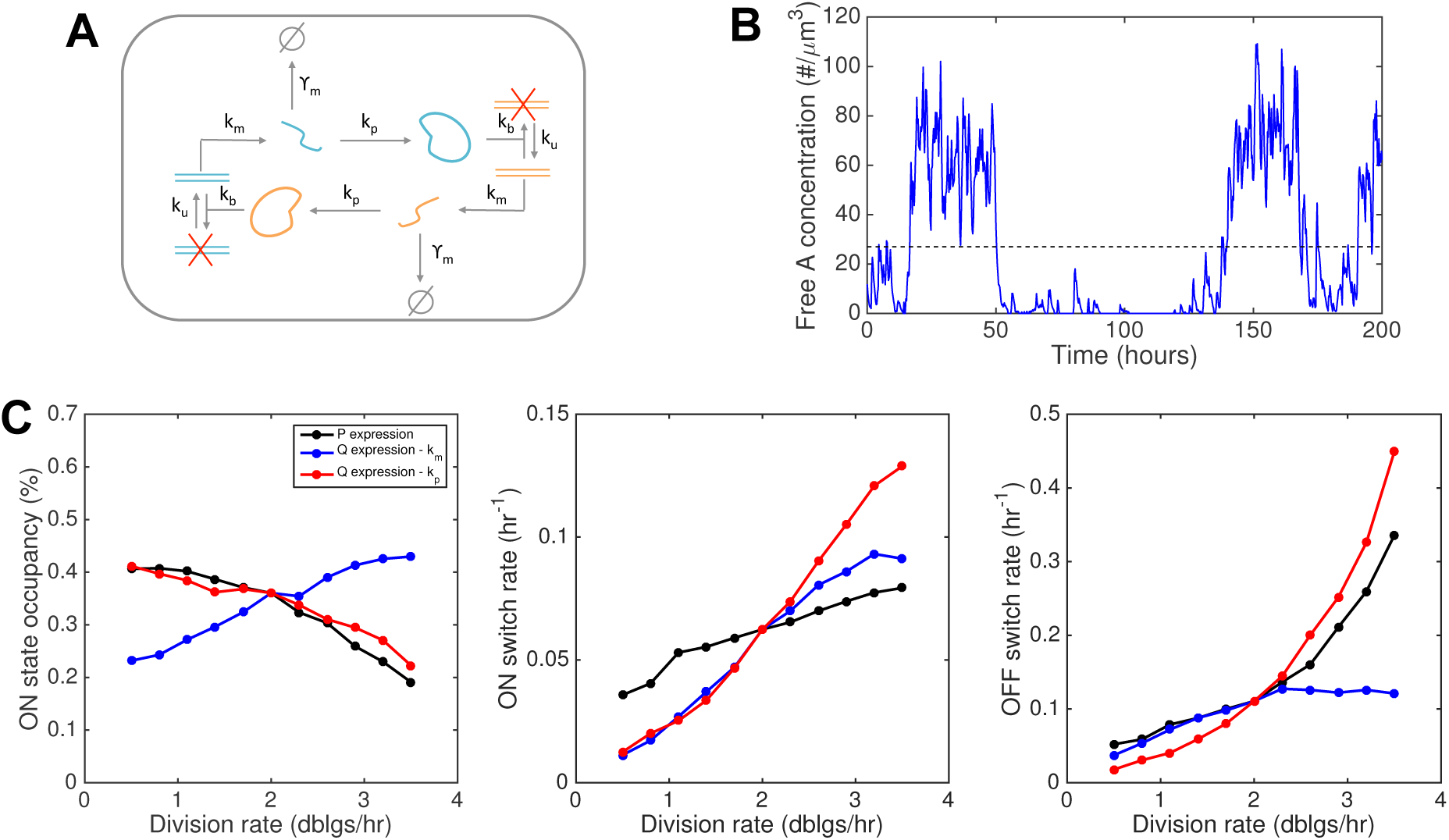
Behaviour of the toggle switch at different division rates. **(A)**Schematic of the toggle-switch circuit. Two proteins *A* and *B* can transcriptionally repress each other by promoter binding. (B) Example simulation of the toggle-switch circuit functioning in growing and dividing cells, showing stochastic switching between high *(ON)* and low *(OFF)* expression for one protein. The threshold separating the two states (black dashed line) is computed using the overall protein distributions (See Methods). (C) Change of the toggle-switch behaviour, quantified by the average time spent in the ON state and the switching rates between the two states, as a function of division rate. The black curve corresponds to *P* expression as in Figure 4, the blue and red curves corresponds to constant average expression maintained either transcriptionally or translationally, as in Figure 3-C,D. Note that when the concentration of one protein type is low, the other is not necessarily high. This is why the *ON* state occupancy is not always 50% despite the symmetry between the two proteins.

We found that the circuit could exhibit bistability (Figure 6-B,C) over the considered range of division rates for constitutive (*P*) expression as well as for *Q* expression by transcriptional or translational adjustment. However, in all cases the circuit behaviour strongly depends on the division rate (Figure 6-C), as illustrated by the change in *ON* state occupancy (the circuit is *ON* when one of the two proteins, the reporter, is in the high expression state). Interestingly, the change of behaviour is very different for different modes of *Q* expression: for translational adjustment, the *ON* state occupancy decreases with the division rate (in a fashion very similar to *P* expression). However, an opposite behaviour is observed for *Q* expression via transcriptional adjustment as *ON* state occupancy becomes positively correlated with division rate.

The *ON* state occupancy reflects the balance between stochastic switching in and out of this state. These rates are both dependent on the division rate (Figure 6-C, middle and right plots). We find that the switching rates increase with the division rate that could suggest random partitioning of mRNA and protein molecules, which is more frequent at high division rates, favours switching as also reported in another study (Lloyd-Price *et al*, 2014). In addition, the observation that at fast growth the *OFF* → *ON* rate rises the most sharply for *Q* expression via translational adjustment is consistent with the high level of protein noise for this mode of regulation at fast division rates (Figure 3-B).

### When gene expression feedbacks on growth: the case of toxin-mediated growth inhibition

So far, the circuits we have considered respond to changes in division rate but they don’t impact cell physiology and growth. However, many natural circuits and some synthetic circuits do influence cell physiology, for example by regulating cell metabolism or cell cycle progression. Even when synthetic circuits are not designed to impact cell physiology, they often do by competing with core cellular processes for global cellular resources, and this has become a major concern for synthetic biologists (Ceroni *et al*, 2015).

In prokaryotes, well-known examples of gene expression feeding back on growth are toxin-antitoxin systems. These systems are involved in bacterial persistence, where a very small subpopulation of slow growing cells naturally arises among a normally growing population. While several existing models represent both the toxin and the antitoxin (Gelens *et al*, 2013, Cataudella *et al* (2013), Fasani & Savageau (2013)), a minimal model where a single protein is toxic for growth is in itself sufficient to generate growth bistability (Klumpp *et al* (2009), Tan *et al* (2009), Rocco *et al* (2013), and Figure 7-B). Here we investigate the behaviour of this kind of model (Figure 7) when both the maximal growth rate reached by a toxin-free cell and the dependency of the transcription rate with the cell growth rate are varied.

**Figure 7:**
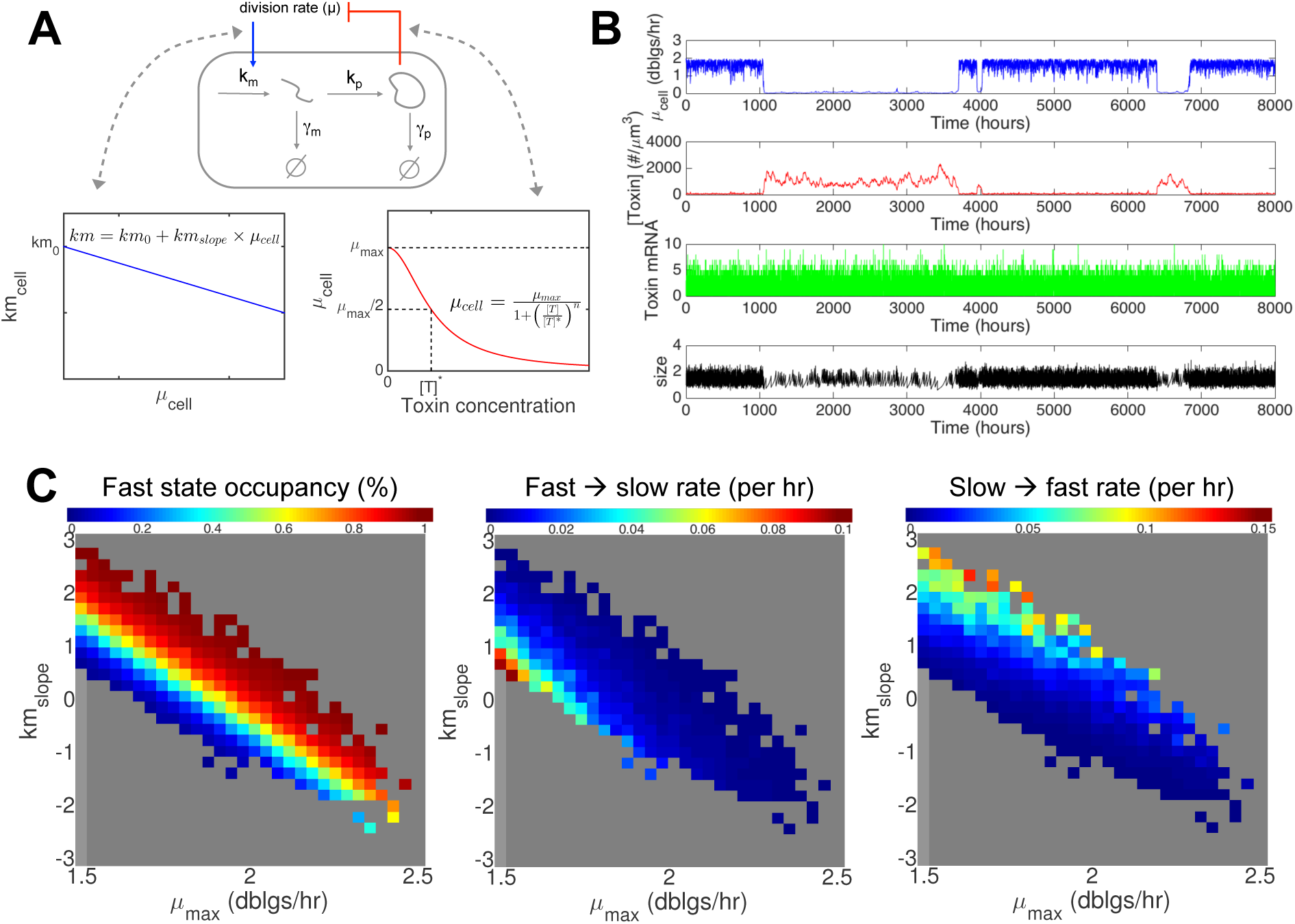
Growth bistability caused by expression of a toxic protein. **(A)**Model description. The instantaneous cell growth rate, which here we assume to be a decreasing function of the expressed protein concentration. In turn, changes in cell growth rate impacts gene expression via the transcription rate. (B) Growth bistability is possible with realistic parameter values (Methods). In the simulation shown, *km_slope_* = 0, meaning that the positive feedback: toxin *→* slower growth *→* more toxin is only mediated by changes in dilution. (C) Influence of growth conditions (*μ_max_*) and growth rate dependence of transcription *(km_slope_)* on growth bistability. For each parameter set, *km_0_* was also adjusted such that *km_cell_*(2 doublings/hr) = 0.28min^−1^. From corresponding simulations, the existence of bistability was tested and corresponding switching rates were estimated (See Methods).

For each parameter set enabling growth bistability (coloured pixels in Figure 7-C), we computed the occupancy of the fast state (Figure 7-C, left) and the switching rates between the slow and fast states (Figure 7-C, middle and right). The occupancy of the fast growing state decreases when the maximal growth rate decreases (Figure 7-C, moving from right to the left), and this behaviour is independent of the dependency of the toxin transcription rate to the cell division rate (i.e. the value of *km_slope_*). Therefore, the system will naturally respond to less favourable growth conditions by increasing the time spent in the slow state.

## Discussion

In this study, we have used detailed simulations of stochastic gene expression in growing and dividing bacteria to investigate the role of division rate in protein noise and dynamics of genetic networks. Our simulations are constrained by data available for *E. coli* related to division rate regulation of constitutive gene expression (Klumpp *et al*, 2009) and single-cell data related to cell size control (Taheri-Araghi *et al*, 2015). For a constitutively expressed gene, we find that coupling transcription but not translation to division rate results in lower protein noise levels. Interestingly, existing data seem to suggest that global regulation of gene expression with division rate mostly acts at the level of transcription (Keren *et al*, 2013; Gerosa *et al*, 2013; Berthoumieux *et al*, 2013; García-Martínez *et al*, 2016), consistent with the idea that lower noise levels are beneficial at fast growth. However, regulation at the level of translation has also been observed (Borkowski *et al*, 2016), which, coupled to transcriptional regulation, could result in non-trivial interplay in terms of gene expression noise regulation. Importantly, those results are independent of the choice of cell cycle noise model, as they remain valid in the absence of noise in cell division size and cell growth rate, at least for gene expression levels where intrinsic noise has a dominant impact.

An important factor that helps to minimise noise in gene expression at fast division rate is increased cell size. Large cell sizes in growth conditions with fast division rate results in higher overall number of mRNA and protein molecules, and reduce noise in gene expression. This is particularly relevant for the regulation of noise in gene expression for proteins belonging to *P* category (Figure 1) as their concentration go down at high division rates. This prediction is supported by experimental data (Nikolic *et al*, 2013). In this study, single-cell expression data (via flow cytometry) was obtained for three promoter-GFP fusion constructs at various growth conditions (Supplemental Figure 6). Of those three promoters, only mglB (a glucose transporter) behaves like a *P* protein: average expression decreases as the growth rate increases. Despite this decrease, expression noise is also decreased, as predicted by our simulations (Figure 4). To more directly validate our predictions on the impact of cell size on expression noise, it would be interesting to leverage recently obtained MreB mutants (Shi *et al*, 2017), which exhibit a strong change of cell volume without reduction of growth rate. Using a constitutive weak promoter integrated in the genome, it would be possible to test whether noise is decreased independently of average concentration and growth rate, simply because of the change in cell size, as predicted by our study. In fact, increasing cell size has been used before to experimentally reduce gene expression noise (Süel *et al*, 2007). In conclusion, we propose that a possible evolutionary reason for microbial cells to grow bigger at fast growth, together with optimization of the surface-to-volume ratio for instance, is to reduce gene expression noise which could be detrimental to fitness at fast growth (Willis & Huang, 2017; Shahrezaei & Marguerat, 2015). At the mechanistic level, the division rate regulation of cell size could be implemented via the division rate regulation of gene expression for proteins involved in cell size control (Basan *et al*, 2015; Bertaux *et al*, 2016).

We restricted our analysis to *E. coli*, because the dependence of gene expression and cell size with the growth rate has been well studied in this species. Importantly, our modeling approach is generic and can be applied to other microbial species. For example, recent data on the dependence of gene expression in *B. subtilis* has been obtained (Borkowski *et al*, 2016), which together with size data (Taheri-Araghi *et al*, 2015) could support predictions about the growth rate dependence of gene expression noise in another species. Interestingly, a recent study (Nordholt *et al*, 2017) has measured an increase in gene expression noise with the growth rate in *B. subtilis* for *P*-like expression, which is different from what has been predicted here for *E. coli* (Figure 4). A potential explanation is that for the growth rate range considered, cell size increase is not strong enough to compensate for the decrease in protein concentration.

In our study, we have modelled extrinsic noise sources such as variability in cell size and cellular growth rate, but we have ignored other sources of noise, such as DNA replication or variability in the abundance of transcription and translation factors (Elowitz *et al*, 2002). The recent study in *B. subtilis* suggests that overall this form of extrinsic noise does not strongly depend on the division rate (Nordholt *et al*, 2017). In addition, the contribution of the DNA replication to protein concentration noise has been found experimentally to be very small (Walker *et al*, 2016).

Our simulations included physiologically relevant levels of partitioning noise, size variability and growth variability. Overall, we observe that the contribution of these factors to protein noise is small (again, intrinsic noise is dominating for our reference gene expression parameters), but that it tends to vary with the division rate for the different cases considered. We also observed the noise in molecular numbers and concentrations do not always behave similarly, as the later directly depends on cell volume. Interestingly, we find that if transcription rate scales with cell size as recently reported in eukaryotes (Padovan-Merhar *et al*, 2015; Kempe *et al*, 2015), the concentration noise becomes independent of noise in cell size control mechanism. In bacteria, there has not been a careful investigation of transcription scaling with cell size and in the absence of such reports we have assumed cell size independent reaction propensities throughout this study.

We then tested how dynamics of simple biochemical networks respond to division rate. As shown by the seminal work of Klumpp *et al* (2009), we find overall that division rate regulation of concentration of *P* proteins can change the average behaviour of biochemical networks significantly. But, as discussed below, we find that even when proteins in the network have a *Q* regulation, the changes in noise properties of the individual gene expression can significantly alter the mean and noise properties of the system.

In the case of a genetic oscillator, we find changes in gene expression and cell size with the division rate can impact the behaviour of oscillatory circuits in a non-trivial manner. Namely, large changes of average expression with the division rate for constitutive expression (*P*) of circuit components render circuit behaviour sensitive to the division rate. However, maintaining constant expression of circuit components (for example via transcriptional or translational adjustment) does not guarantee full robustness of circuit behaviour against changes in division rate. Robustness might require more complex, circuit-specific dependencies of gene expression with the division rate, or even specific circuit architecture (Paijmans *et al*, 2016). Interestingly, we observed a ‘U’ shape dependency of noise on division rate suggesting that there could be an optimally robust growth condition for a specific network design and parameter combination, which is relevant to appropriate function of natural biochemical systems or synthetic systems.

The toggle switch circuit behavior is strongly dependent on the division rate and on the type of gene expression dependency with the division rate. So, this suggests the simple toggle switch circuit is not going to perform robustly across growth conditions. As for the oscillator circuit, maintaining average expression is not sufficient to generate a division rate independent behaviour. Moreover, this example shows that even when average expression is maintained, whether it is maintained via adjustment of transcription or translation matters, as the circuit behaves differently in either situation. Those predictions can be tested experimentally, for example using a recent implementation of the circuit, for which a detailed stochastic model (validated for a single growth condition) is available (Lugagne *et al*, 2017).

In the case of simple models of persistence induced by the expression of a toxic protein in single growing and dividing cells, we could investigate the impact of growth conditions and gene expression dependency with the cell growth rate on the emergence of growth bistability. The role of growth conditions in prevalence of persister cells is a very relevant problem as the growth conditions of bacteria during infection are likely to be altered by the immune system and therapeutic treatments for instance. We found that the persister state is favoured in poor growth conditions. A similar observation has been made in another theoretical study with a more detailed model of toxin and antitoxin interactions (Gelens *et al*, 2013). To further validate those modelling results, it would be interesting to experimentally assess if and how growth conditions regulate the probability of the non-growing persistence phenotype.

In molecular systems biology, we use models of biochemical networks to validate our mechanistic understanding of the system under study. We propose that such models should be tested also against data collected across cellular division rates. If the behaviour of the system is observed to be robust to growth conditions, then our models should be able to capture this robustness. Conversely, describing the ways in which the system behaviour changes across growth conditions is key to refine our models and therefore our mechanistic understanding of the system under study.

In synthetic biology, we often desire to build a system that either functions robustly at a particular growth condition or across a range of growth conditions. Our study shows that stochastic models of synthetic biochemical networks in growing and dividing cells coupled with data on the regulation of gene expression and cell size across division rates are essential to optimal design of system topologies that achieve robustness against changes in cellular division rates.

## Data availability

The code used for all simulations is available on GitHub (https://github.com/ImperialCollegeLondon/coli-noise-and-growth).

## Author’s contributions

F.B., S.M. and V.S. designed the study. F.B. wrote the code and performed simulations. F.B. and V.S. analyzed and interpreted the simulation data. F.B., S.M. and V.S. wrote the manuscript. All authors gave final approval for publication.

## Acknowledgments

We would like to thank Philipp Thomas and Marc Sturrock for feedback on our manuscript and members of the Marguerat and Shahrezaei groups for discussions. We thank Suckjoon Jun for sharing the *E. coli* size data across growth conditions. We acknowledge Matthew Robb for his preliminary work on this project during his PhD. This work is supported by a Leverhulme Research Project Grant (RPG-2014-408) awarded to S.M. and V.S. S.M. is supported by the UK Medical Research Council.

## Methods

### Basic model for gene expression in growing and dividing cells

We describe first our basic model for gene expression in growing and dividing cells. mRNA molecules are randomly synthetized and degraded at rate *k*_*m*_ and *γ*_*m*_ respectively. Stochastic synthesis of protein from each mRNA occurs at rate *k_p_*. Protein molecules are assumed to be stable (except for *A* in the oscillator circuit). Cell volume is growing exponentially at a fixed rate between *V*_*birth*_ and *V*_*div*_ = 2 *V*_*birth*_, then cell division is triggered (for the case including cell size control and variability see below). At cell division, molecules are randomly split between daughter cells and the volume is halved. In simulations, only one of the daughter cell is considered for further simulation (hence mimicking the ‘mother machine’ microfluidic experiments for a symmetrically dividing cell (Wang *et al*, 2010). We note that as expected, in our simulations, partitioning noise at division reduces the correlation between mRNA and protein levels (from 0.3 to 0.2 with the reference gene expression parameters), consistently with experimental observations (Golding *et al*, 2005).

### Reference gene expression parameters

Realistic (Taniguchi *et al*, 2010) parameters for *E. coli* gene expression have been used (*k*_*m*_ = 0.28*min*^−1^, *γ*_*m*_ = 0.14*min*^−1^, *k*_*p*_ = 0.94*min*^−1^, *μ* = 2 doublings/hr). This corresponds to an mRNA half-life of 5 min, an average mRNA number at birth of 1 molecule and an average protein number at birth of 50 molecules.

### Quantification of protein noise

Throughout our study, noise is quantified by using coefficient of variability (CV), which is defined as standard deviation divided by the mean. Except when stated otherwise, we call protein noise the noise in protein *concentration*, which is physiologically more relevant than molecule numbers. Also, noise is computed among newly born cells in order to eliminate cell cycle stage contributions. Similar trends are observed in the middle of the cell cycle.

### Realistic modelling of cellular growth rate and cell size variability across growth conditions with noisy linear maps

We use *noisy linear maps* (Tanouchi *et al*, 2015, see Supplemental Figure 1 for a description of our model with NLM) with parameters inferred from mother machine data in different growth conditions (Taheri-Araghi *et al*, 2015). The data contains around 100K cell cycles per condition. *a* is estimated by linear regression of *l*_*div*_ (division length) vs *l*_*birth*_ (birth length). *σ*_1_ is by definition related to the residual of this regression. *σ*_2_ is estimated from the variance of 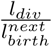 where 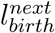 is the birth length recorded just after the division at *l*_*div*_. Once *a* has been estimated, *b* is chosen such that the model predicted average birth size 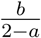 matches the average birth size observed in the data.

### Impact of size-dependent transcription

In all our models, the reaction propensities for transcription, translation and mRNA decay are independent of cell volume. However, transcription rates have been recently reported in eukaryotes to depend on size (Padovan-Merhar *et al*, 2015; Kempe *et al*, 2015). Because the dependency of protein noise on NLM parameters could be affected by size-dependent transcription, we have repeated the analysis presented in Figure 2 using transcription rates proportional to cell size (Supplemental Figure 7). Interestingly, we find that in this case the protein concentration noise is reduced and becomes independent of the NLM parameters. We obtain very similar results if we assume translation rate is size-dependent (not shown).

### Modelling *Q* expression by transcriptional or translational adjustment

For a stable protein, it is possible to derive an analytical expression for the average number of protein molecules at birth: 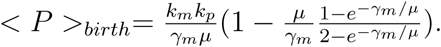

This expression was used to compute the transcription or translation rate achieving a given average protein concentration (Figure 3 and Supplemental Figure 4). The fact that our simulation algorithm produced the expected dependency of average protein concentration validates its implementation. In the case of active protein degradation (as for *A* for the oscillator circuit studied in Figure 5), we used simulations and the MATLAB scalar optimization function *fminsearch* to compute the transcriptional or translational rate adjustment enabling to maintain a constant average concentration at birth.

### Modelling *P* expression

For Figure 4, we have used division rate dependencies of gene expression parameters from (Klumpp *et al*, 2009) (cf Supplemental Figure 4). For modelling *P* expression in the oscillator circuit (Figure 5), for simplicity we only used the effective transcription rate (and cell size) division rate dependency, as change in translation rate per mRNA or mRNA degradation rate are small.

### Oscillator circuit

The model structure and parameterization is adapted from (Vilar *et al*, 2002). The *Act* protein can transcriptionally activate its own expression as well as the expression of another protein *Rep* by promoter binding. *Act* is short-lived while *Rep* is stable. *Act* and *Rep* can form a complex. The same model reactions were used, but we also explicitly model growth and division (including random partitioning of free *Act* and free *Rep*, but we do not model gene replication and consider a single copy of each promoter which is always inherited by daughter cells). The volume dependency of bi-molecular reactions is also accounted for. As reference parameters (i.e. corresponding to an intermediate *E. coli* division rate of 2 doublings per hour, at which optimal circuit behavior should be obtained), we used the same parameters as Vilar and colleagues, except that the *Rep* degradation rate was set to 0 (the original value, corresponding to a ~200 min half-life, was accounting for dilution only), the active degradation rate of *Act* was scaled up to maintain a constant ratio with the division rate, the *Rep* translation rate was scaled up by the same factor, and all transcription rates were scaled by this factor (~7).

The resulting values are:

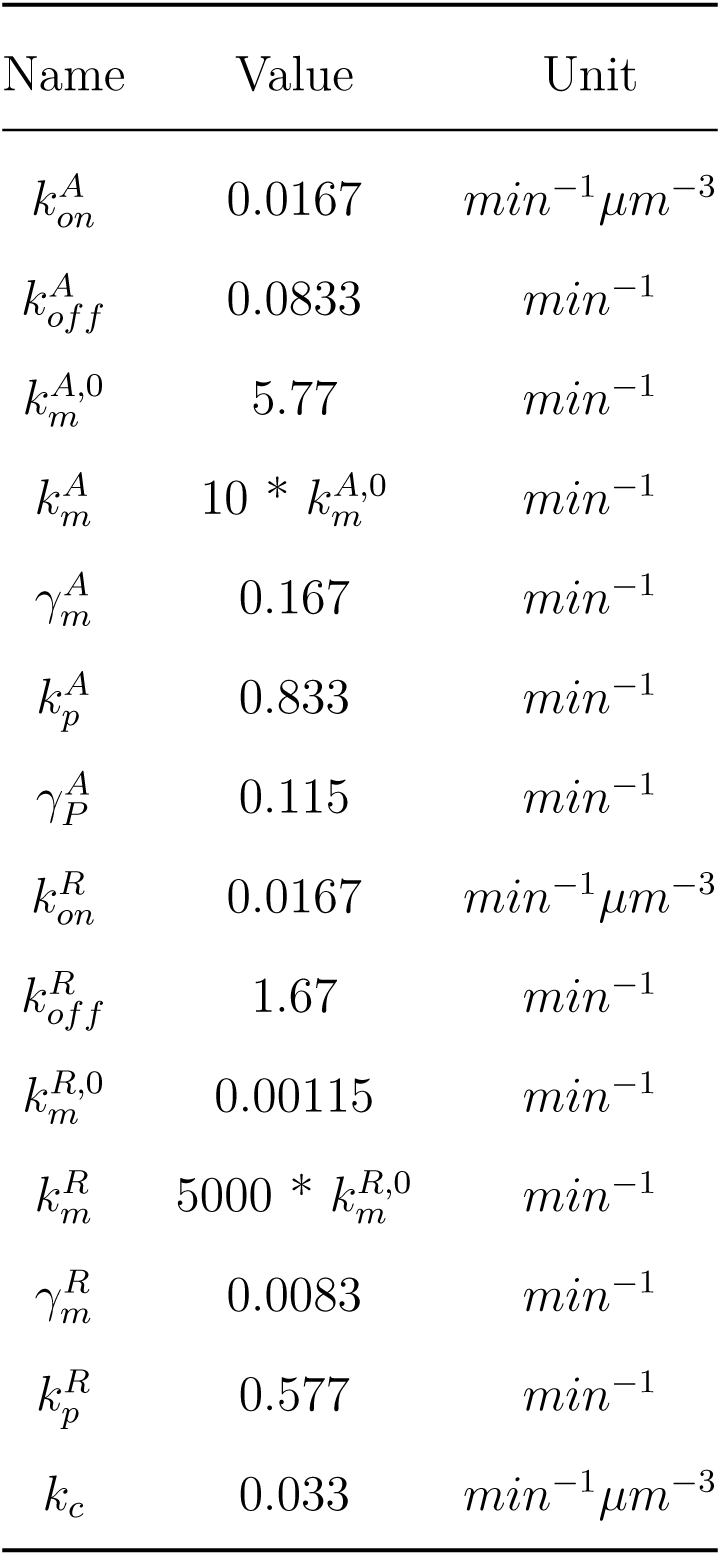

| Name | Value | Unit |
| --- | --- | --- |
| $k_{on}^A$ | 0.0167 | $min^{-1}\mu m^{-3}$ |
| $k_{off}^A$ | 0.0833 | $min^{-1}$ |
| $k_m^{A,0}$ | 5.77 | $min^{-1}$ |
| $k_m^A$ | $10 * k_m^{A,0}$ | $min^{-1}$ |
| $\gamma_m^A$ | 0.167 | $min^{-1}$ |
| $k_p^A$ | 0.833 | $min^{-1}$ |
| $\gamma_P^A$ | 0.115 | $min^{-1}$ |
| $k_{on}^R$ | 0.0167 | $min^{-1}\mu m^{-3}$ |
| $k_{off}^R$ | 1.67 | $min^{-1}$ |
| $k_m^{R,0}$ | 0.00115 | $min^{-1}$ |
| $k_m^R$ | $5000 * k_m^{R,0}$ | $min^{-1}$ |
| $\gamma_m^R$ | 0.0083 | $min^{-1}$ |
| $k_p^R$ | 0.577 | $min^{-1}$ |
| $k_c$ | 0.033 | $min^{-1}\mu m^{-3}$ |

To compute the period and amplitude of oscillations in free *R* concentration, we used the MATLAB function *findpeaks* on very long (200K minutes) mother machine traces, requiring a minimum peak amplitude of 25% of the maximum value in the trace. We verified visually the behavior of the peak detection algorithm for each simulation.

### Toggle switch circuit

The model structure and parameters are completely symmetric for the two proteins repressing each other. There is no cooperativity in the repression, as it is not required to obtain stochastic switching, consistently with (Lipshtat *et al*, 2006). As for the oscillator circuit, the volume dependency of bi-molecular reactions (only promoter binding here) was accounted for. We assumed that transcription is completely blocked when the promoters are bound, and that the promoter binding and unbinding rates are independent of the division rate.

The reference parameter values are:

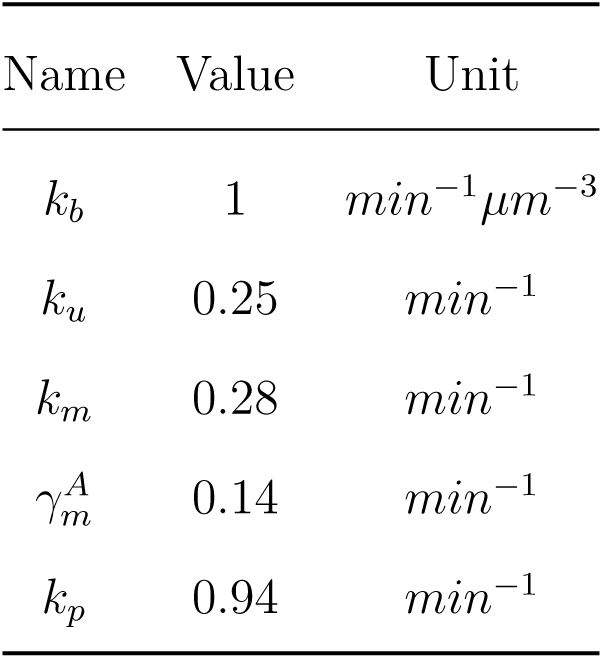

Detection of bistability (always the case for simulations shown in Figure 5), threshold identification and computation of switching rates were performed as follows. A very long (500 thousands hours of biological time) single-lineage trace (one output every 15 minutes) of the free *A* concentration is obtained by simulation. This trace is then discretized into 50 equal size bins from zero to the maximal value of the trace. The following algorithm is then applied on this discretized distribution: *(1)* identify the highest mode (i.e. the most populated bin); *(2)* iteratively identify next highest mode and ask whether they are corresponding to a neighbor bin of the highest mode (then it is not the second mode of a bimodal distribution) OR if there exists populated, lower height bins in-between (indicative of bimodality); *(3)* in the latter case, to avoid incorrect detection of bimodality because of finite sampling of the distribution, the secondary mode is required to be more than 5% of what an uniform distribution would give.

### Growth bistability caused by expression of a toxic protein

As previously, stochastic gene expression of a protein is simulated in growing and dividing cells. However, the protein is a toxin inhibiting cell growth: the instantaneous growth rate of the cell *μ*_*cell*_ is a decreasing Hill function of the toxin concentration (hence it is not anymore constant during the cell cycle). Also, the impact of growth conditions is not modeled anymore with condition-specific noisy linear maps, as they are not adapted to situations with very heterogeneous growth rates between cells in a given condition. We rather use a parameter *μ*_*max*_ representing the toxin-free cellular growth rate. For simplicity, to model cell division size and its variability we use a single noisy linear map across growth conditions. Finally, to represent the dependency of gene expression with the cell growth rate, we assume that the toxin transcription rate is a linear function of *μ*_*cell*_. The reference parameter values are:

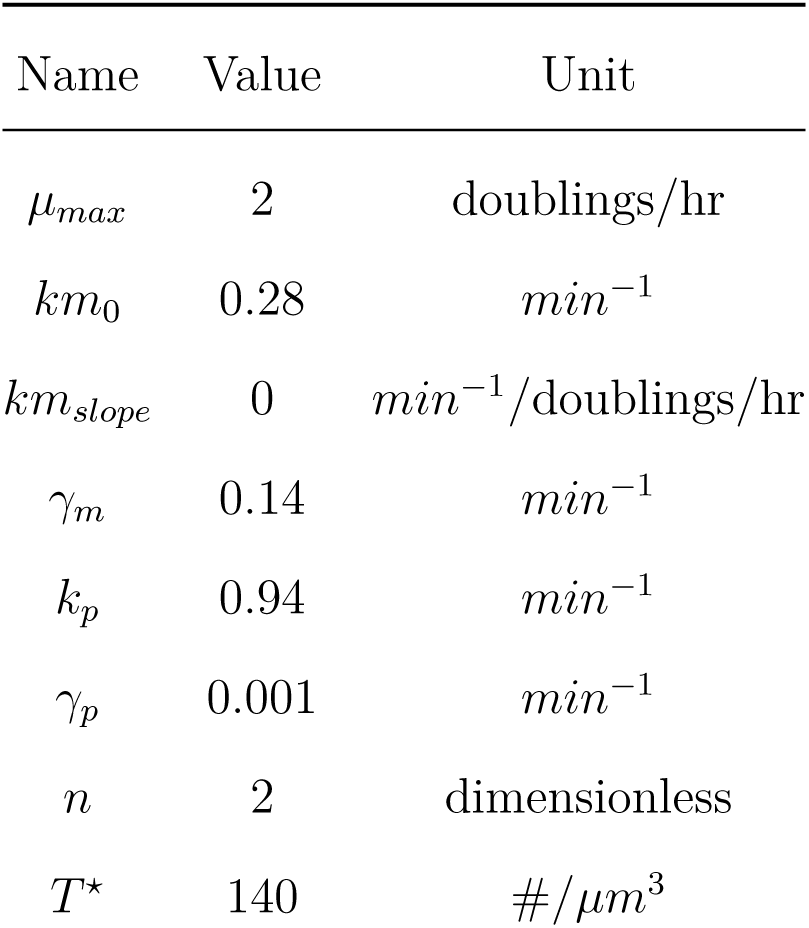

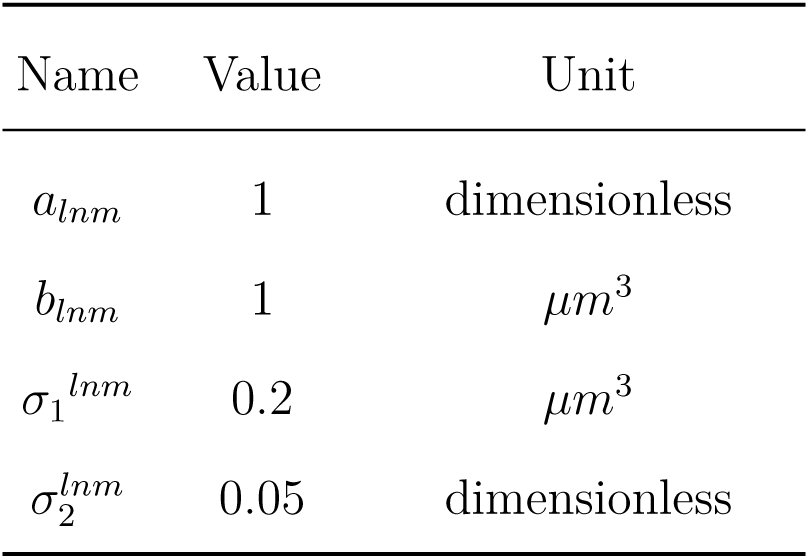

Note that because *km*_*slope*_ = 0, the positive feedback toxin → growth slow down → more toxin is only mediated by a change of dilution (as in (Rocco *et al*, 2013)). Also note that it is necessary to assume that protein degradation is non-zero to allow bistability, as otherwise exit of the slow state is impossible.

For Figure 7-C, for each parameter set, the existence of bistability, threshold identification and switching rates computation for the instantaneous cell growth rate *μ*_*cell*_ were performed as for the toggle switch circuit analysis (except that simulation duration for each single-lineage trace was 60 thousands hours of biological time, with one output every 10 minutes, and the number of bins used was 20).

Grey indicates parameter sets for which the lineage simulation of 60 thousands hours (~120 thousands generations) either did not lead to a bimodal distribution of *μ*_*cell*_, or did lead to such bimodal distribution, but with less than 10 switches fast → slow → fast, preventing an accurate estimate of switching rates in reasonable computational time.

### Simulation algorithm

We describe here the general simulation algorithm used for all models. Between fixed timesteps (6 seconds), cell volume is considered constant, and the Gillespie algorithm is used to simulate stochastic molecular reactions (more sophisticated simulation methods exist (Lu *et al*, 2004; Shahrezaei *et al*, 2008), but this one is simple to implement and accurate as long as the timestep is small enough). Then, the cell volume is updated according to the instantaneous exponential growth rate, it is checked whether cell division should occur, and if so, cell division and molecules partitioning is realized.

